# Harnessing mRNA for the expression of monoclonal IgG and IgA in non-human primates

**DOI:** 10.1101/2025.04.09.648023

**Authors:** Romy Rouzeau, Hayden R. Schmidt, Cailin Deal, Joel D. Allen, Dawn M. Dudley, Iszac Burton, Eva G. Rakasz, Obadiah Plante, Max Crispin, Andrea Carfi, Devin Sok

**Author notes:** **Corresponding authors contact information:** Andrea Carfi, Devin Sok. Denotes equal contribution (Romy Rouzeau, Hayden R. Schmidt, Cailin Deal).

## Abstract

Monoclonal antibodies (mAbs) are an increasingly essential class of medicines across many disease areas (1). In the human body, there are five antibody isotypes, each with potential prophylactic or therapeutic benefits for different disease indications. However, 97% of all clinically approved mAbs are produced as the IgG isotype, partly due to differences in half-life, but largely due to challenges associated with recombinantly producing non-IgG isotypes like IgM or IgA, which have additional N-linked glycan sites and can present as multivalent oligomers. One potential solution to this challenge is to express mAbs *in situ* using mRNA encapsulated in lipid nanoparticles (LNP), bypassing the need for recombinant protein production (2).

Here, we demonstrate the feasibility of expressing a mAb as both IgG and IgA in non-human primates (NHPs) using mRNA-LNPs. We express ePGDM1400v9, a broadly neutralizing mAb targeting human immunodeficiency virus (HIV), in both IgG1 and IgA2 formats by infusing NHPs with LNPs containing the appropriate mRNA. Though IgG1 expression levels were higher than those of IgA2, both formats were detectable in serum within one day of LNP infusion in all NHPs, and both were detectable in mucosal secretions of most animals. Importantly, serum mRNA-produced IgG1 and IgA2 retained HIV-neutralizing function. Furthermore, mass spectrometry analysis confirmed that mAbs of either isotype produced *in situ* exhibited glycosylation patterns highly similar to that of native antibody, which is likely to confer therapeutic advantages. Altogether, this work demonstrates the feasibility of using mRNA-LNPs to express native-like mAbs of non-IgG isotypes in primates and enables further development of non-IgG mAb constructs.

**Significance:** Monoclonal antibodies (mAbs) are a rapidly growing class of essential medicines across diverse disease areas and applications. However, the impact potential of mAbs is limited by challenges in production and purification, which favors the use of the IgG antibody isotype despite the disease-specific advantages that other isotypes might offer. One way to overcome this challenge is to express mAbs *in situ* using messenger ribonucleic acid delivered by lipid nanoparticles (mRNA-LNPs). Here, we show that an anti-human immunodeficiency virus (HIV) mAb, ePGDM1400v9, can be expressed as two different antibody isotypes, IgG and IgA, in nonhuman primates (NHPs) by mRNA-LNP delivery. We demonstrate that both mAb isotypes retain function and exhibit native-like glycosylation patterns that are not achievable with conventional recombinant mAbs.

## Introduction

Monoclonal antibodies (mAbs) are an essential class of therapeutics for indications across cancer, neurological disease, autoimmune disorders, and infectious disease (1). These biologics hold several advantages over traditional small molecule drugs, including their target specificity and versatility, excellent safety profile, amenability to engineering, and (as demonstrated during the SARS-CoV-v2 pandemic) the rapid speed with which they can be identified and developed. These properties have made mAbs one of the fastest growing classes of medicines over the decade (1).

In nature, there are five main isotypes of antibodies that differ in their constant domains, all exhibiting distinct immunological effector functions and recruitment of different immune effector cells. Despite this immunological diversity, almost all mAbs that have been approved or are pending approval are of the IgG isotype (97%) due to the ease of manufacturing and favorable pharmacokinetic properties. As such, less research and effort has been made on optimizing production and purification of alternative isotypes such as IgM and IgA, even when these isotypes may provide disease-specific therapeutic advantages over IgGs.

A potential solution to these challenges is to express mAbs *in situ* using lipid nanoparticle (LNP)-encapsulated messenger ribonucleic acid (mRNA) (3). As a platform, mRNA inherently bypasses the need for protein expression and purification (2), thereby enabling characterization and development of molecules such as IgA, which has been largely untapped because of manufacturing difficulties. The chemistry and manufacturing of mRNA remains the same, regardless of protein, thus allowing for rapid production of different molecules. An additional advantage over recombinantly produced mAbs is the potential for more native-like glycosylation patterns when produced *in situ* from mRNA (4). This has proven to be advantageous for complex molecules like IgA as a lack of sufficient sialylation, which is reduced in common antibody production cell lines, can result in rapid clearance *in vivo* (5). Nonetheless, IgA mAbs could be transformative therapeutics against mucosal pathogens due to their privileged mucosal access as IgA dimers, complexed together with the J chain (5, 6), which enables transcyotosis via the polymeric immunoglobulin receptor (pIgR).

To date, mRNA-delivered mAbs against human immunodeficiency virus (HIV) (7, 8), tumors (9, 10), botulin toxin (10), rabies virus (10), respiratory syncytial virus (11), and Chikungunya virus (12) have been shown to be protective in mice. The mRNA-encoded anti-Chikungunya virus mAb (mRNA-1944) expressed to theoretical protection levels in both non-human primates (NHP) (12) and healthy adults in a Phase I clinical trial (13). The above mAbs represent various formats and constructs, though all are expressed as IgGs or in formats that have been validated in the clinic using recombinant protein products. More recently, we reported the successful expression of IgA mAbs in mice, which were able to prevent mucosal colonization of pathogenic *Salmonella* and *Pseudomonas* (4).

Here, we build upon this work by demonstrating that ePGDM1400v9, a broadly neutralizing antibody (bnAb) against HIV (14), can be expressed in NHPs as either a human IgG1 or a human IgA2 mAb. We show that mRNA-expressed ePGDM1400v9 persist in serum for time periods comparable to either native IgA2 or recombinant IgG1 (with half-life extension through a “LS” mutation (15)), and that these mAbs are capable of neutralizing HIV pseudovirus *in vitro*. Additionally, glycan analysis of ePGDM1400v9 IgG1 and IgA2 produced in situ reveals native-like host-specific glycosylation, with the attachment of N-glycolylneuraminic acid, which is a modification not commonly observed in CHO or HEK293F cells used for recombinant production. Altogether, this work demonstrates that mAbs can be produced in situ in primates via mRNA-LNPs as either IgG or IgA isotypes, and acquire native-like glycosylation while retaining the same functional activities as their recombinant counterparts.

## Results

### Generation and validation of antibodies and assays

To facilitate the accurate measurement of ePGDM1400v9 in NHP tissues and serum, we acquired anti-idiotype antibodies specific to the variable domain of ePGDM1400v9 through a contract research organization (see methods for details). The anti-idiotype antibody used in this work, aPGDM01, was produced recombinantly as a mouse IgG2a and was shown to bind to recombinant ePGDM1400v9 as a Fab, human IgG1, human IgG2, monomeric human IgA2 and dimeric human IgA2 with high affinity and specificity by surface plasmon resonance (Figure S1A). Importantly, though recombinant ePGDM1400v9 IgA2 was purified as a mixture of monomers and dimers (Figure S1B), ELISAs using separated dimer or monomer peaks confirmed that aPGDM01 binding was not influenced by ePGDM1400v9 IgA2’s oligomeric state (Figure S1C). To measure ePGDM1400v9 concentrations in NHP sera, an electrochemiluminescence (ECL) assay was developed. This assay was able to accurately measure recombinant ePGDM1400v9 spiked into NHP serum for both IgG1 (Figure S1D) and IgA2 (Figure S1E) formats.

### mRNA-derived ePGDM1400v9 is produced *in vivo* and is detectable in serum and mucosal secretions

We sought to determine the kinetics and quantity of expression of mRNA expressed ePGDM1400v9 as an IgG1 and an IgA2 (Figure 1). Two groups of 4 rhesus macaques (8 total) received a 1 mg/kg intravenous transfusion of mRNA-LNPs encoding for either ePGDM1400v9 human IgG1-LS (the “LS” mutation conferring improved half-life (15)) or ePGDM1400v9 IgA2 (Figure S2). On day 4 post-transfusion, two animals from each group were necropsied, and select tissues were harvested for biodistribution analysis. The remaining animals were necropsied on day 70 post-transfusion and the same tissues were harvested. No adverse events were observed in any of the animals during the study.

**Fig. 1:**
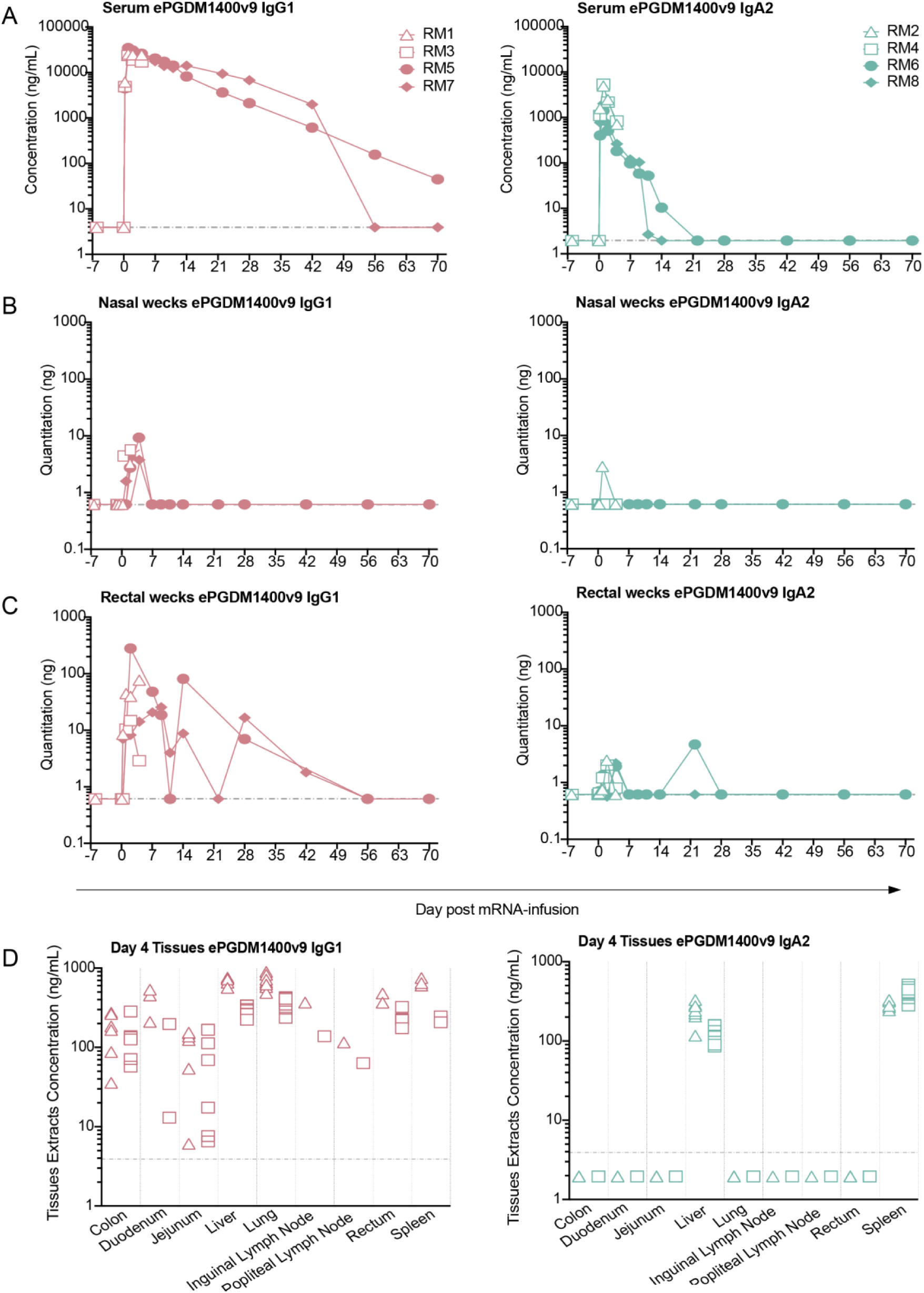
mRNA-mediated expression ePGDM1400v9 IgG1 and IgA2 in NHP serum and tissues. Longitudinal quantification of ePGDM1400v9 IgG1 or IgA2 in mRNA-transfused macaques in: sera (A) by ECL, nasal (B) and rectal (C) wecks by ELISA. The nasal weck sample from IgA2-receiver RM2 collected at d2 post-transfusion was excluded due to the presence of blood on the weck. Similarly, rectal weck samples from IgG1-receiver RM5 collected d1, d4, d21 and d42 post-transfusion, and the sample from IgG1-receiver RM7 collected at d70 post-transfusion, were excluded from analysis due to the presence of blood on the weck. Quantification of ePGDM1400v9 IgG1 or IgA2 (D) in tissue in necropsied animals at day4 post-transfusion by ECL. Note that tissues were not perfused at necropsy, so the measurements here represent the total antibody in each tissue and the associated vasculature.

All animals of both groups exhibited expression of ePGDM1400v9 in sera that persisted up to 4 d post-transfusion, when two animals from each group were necropsied (Figure 1A, Table S1). Across all animals, serum levels of both IgG1 and IgA2 ePGDM1400v9 peaked at 1-day post-transfusion, though IgG levels were significantly higher. Among the four NHPs in each group, peak ePGDM1400v9 IgG1 serum levels ranged from 23.7 – 34.5 µg/mL, while peak ePGDM1400v9 IgA2 serum levels ranged from 1.4 – 5.4 µg/mL. Of the two remaining (not necropsied at 4 d post-transfusion) NHPs that received the IgG1 mAb, one exhibited antibody over the limit of detection (LOD) up to 70 d, while the other dropped below the LOD between 42 d and 56 d. Of the two remaining IgA2-receiving NHPs, one dropped below the LOD between 10 d and 14 d, and the other between 14 d and 21 d.

We also measured total ePGDM1400v9 in rectal (Figure 1B) and nasal (Figure 1C) Weck-cel sponges (wecks) by ELISA to measure mRNA-derived antibody at mucosal surfaces. In rectal wecks, ePGDM1400v9 IgG1 was detectable in all animals at low levels up to 4 d post-transfusion, while ePGDM1400v9 IgA2 was detectable in most but not all animals over the same period. No ePGDM1400v9 was detectable in any rectal swabs after 4 d post-transfusion. The amount of ePGDM1400v9 IgG1 detected in nasal swabs was much lower than in rectal wecks, and while all animals exhibited detectable antibodies up to 4 d post-transfusion, not all animals had detectable antibody at all timepoints. IgA2 was only detected from one nasal weck from one animal at 1 d post-transfusion. Note that for both rectal and nasal wecks, timepoints were excluded for some animals if blood was observed on the weck (see figure legend for details). Altogether, these data suggest that a small proportion of mRNA-derived ePGDM1400v9 IgG1 and IgA2 did reach mucosal surfaces. Expressing the mAb as an IgA2 did not confer any detectable benefit with respect to mucosal trafficking, though this may be due to limitations in detection methods used and experimental design (see discussion).

### Biodistribution of mRNA-derived ePGDM1400v9 four days post-transfusion

We also measured mRNA-delivered ePGDM1400v9 in select tissues from animals sacrificed at 4 d post-transfusion (two animals per group). When expressed an IgG1, ePGDM1400v9 was detectable in all tissues tested (Figure 1D, Table S1). However, when expressed as an IgA2, the mAb was only detectable in the liver and spleen, where mRNA-LNPs are known to accumulate and express their targets (16). These data suggest that while expression levels achieved for the IgG1 construct were high enough to achieve systemic distribution, the IgA2 construct was not detectable in most tissues at 4 d post-transfusion. Note that the tissues examined were not perfused, so ePGDM1400v9 levels measured here represent the total amount of antibody in both the tissue itself and the associated vasculature.

### ePGDM1400v9 IgG and IgA from transfused macaques neutralizes HIV pseudovirus *in vitro*

We next sought to verify that the ePGDM1400v9 mAbs produced in the primates by mRNA retained expected neutralization activity against sensitive HIV strains. Both plasma and sera from transfused animals were screened against a panel of three different HIV pseudoviruses, each with a different degree of sensitivity to ePGDM1400v9 IgG1 (Figure 2A). Consistent with the observed expression of antibody in all animals, sera were able to selectively neutralize sensitive pseudoviruses (Figure 2B, 2C, 2D and Table S2A). No neutralization was observed against the negative control MLV, at any time point (data not shown). Importantly, the serum ID_50_ for each sample correlated well with the serum ePGDM1400v9 concentration measured by ECL (Figure 2D). Therefore, we used the ePGDM1400v9 serum concentrations determined by ECL and the neutralization assay ID_50_s to calculate estimated IC_50_ values for each group against each virus, which facilitated a direct comparison to recombinant ePGDM1400v9 IgG1 (Figure 2E). We observed that the potency of the ePGDM1400v9 IgG1 and IgA2 against sensitive pseudoviruses (T250-4 and CE1176) was statistically indistinguishable from recombinant ePGDM1400v9 IgG1, suggesting that the mAbs produced *in situ* by mRNA had similar neutralization activity to the recombinantly produced mAb.

**Fig. 2:**
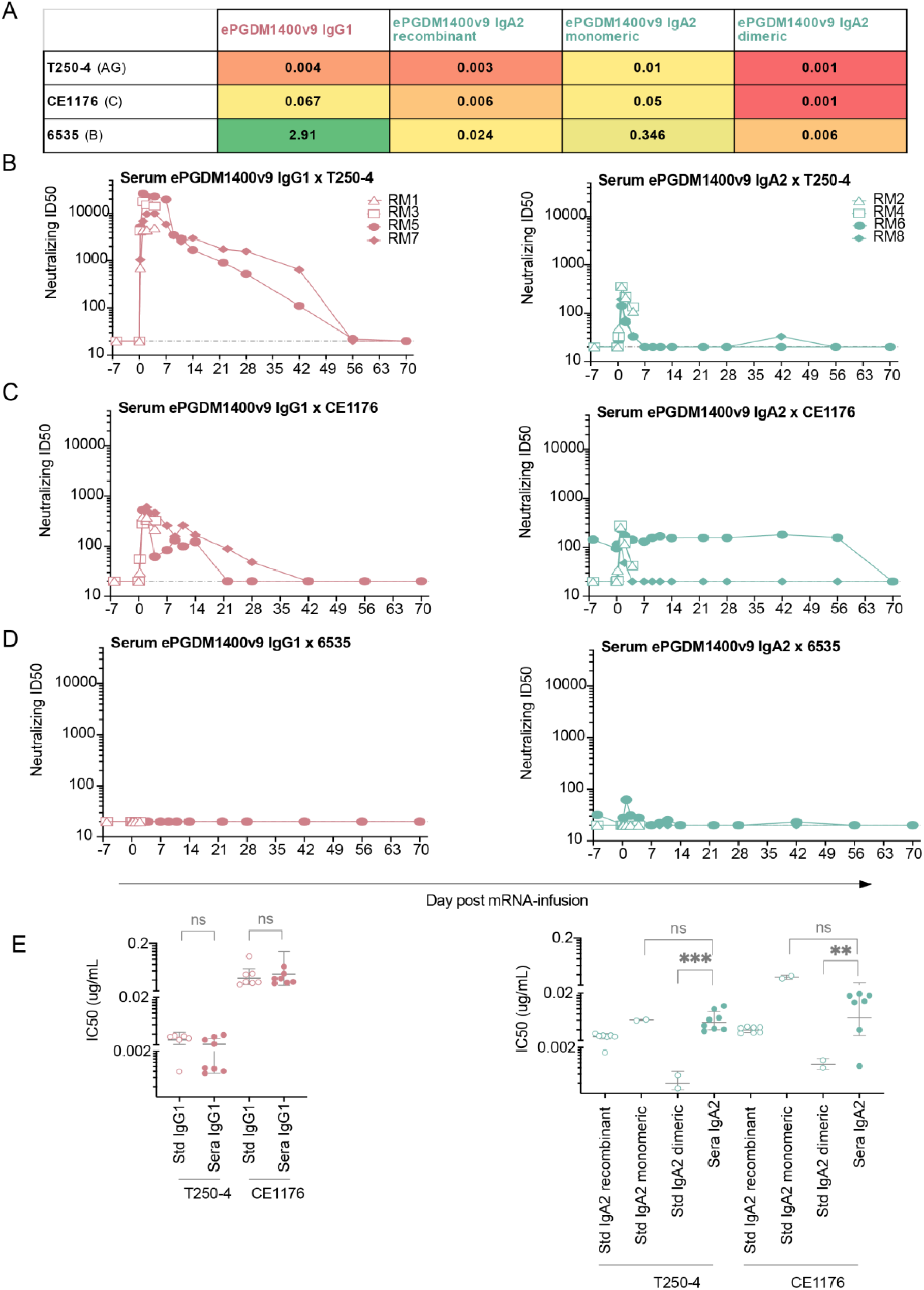
mRNA-derived ePGDM1400v9 IgG1 and IgA2 from NHP sera neutralizing activity as assessed by TZM-bl assay. (A) *in vitro* IC50 of the 3 cross-clade pseudoviruses that were selected based on their low (6535), mid (CE1176) or high (T250-4) neutralization by ePGDM1400v9 IgG1. (B to D) Neutralization ID50 of IgG1 and IgA2 infused sera at each collected timepoint against T250-4 (b), CE1176 (c) and 6535 (d). (E) mRNA-derived ePGDM1400v9 IgG1 (left) and IgA2 (right) IC50 against T250-4 and CE1176 extrapolated based on the neutralization ID50 and the serum concentration evaluated Fig 2B and 2C, and aligned with control isotype IC50 for each of the 3 pseudoviruses.

However, we did note that the *in vitro* neutralization potency of the mRNA-derived IgA2 mAb did not match as closely to the recombinant IgG1 as the mRNA-derived IgG1. Therefore, we repeated these experiments for samples obtained at 1 d and 4 d post-transfusion, ensuring we used recombinant ePGDM1400v9 IgA2 as a comparator. The recombinant IgA2 was produced via transient transfection of ePGDM1400v9’s light chain (kappa), heavy chain (IgA2), and the j-chain required for IgA dimerization which resulted in a mixture of monomer and dimer (Figure S1D). Since dimerized IgA2 is expected to rapidly traffic to the mucosa, the IgA2 in the serum was likely to be nearly 100% monomeric. Therefore, we separated the recombinant monomeric and dimeric species using size exclusion chromatography and compared the estimated IC_50_ neutralization potency of the sera to recombinant mixed, monomeric, and dimeric species (Figure 2E).

We observed that dimeric ePGDM1400v9 IgA2 was more potent than the monomeric form, and that the serum IgA2 was generally more similar to the monomeric species in potency, which is consistent with expectations. Therefore, ePGDM1400v9 produced in NHPs by mRNA transfusion retains *in vitro* neutralization activity as both an IgG1 or an IgA2, and most if not all of the serum IgA2 was monomeric, which is consistent with the canonical biodistribution of monomeric and dimeric IgA.

### Glycosylation analysis of mRNA-derived ePGDM1400v9 IgG1 and IgA2

N-linked glycosylation plays a key role in modulating immunoglobulin effector functions and is heavily dependent on the producer cell from which the antibodies are derived. As mRNA derived antibodies are produced *in situ*, the factors that influence glycosylation such as metabolite availability as well as the expression level and type of glycosyltransferases, will differ between typical CHO cells used to produce therapeutic Ig and the various cell types that take up the mRNA *in vivo*. Human IgA2 contains 4 potential N-linked glycosylation sites defined as an amino acid sequon consisting of NxS/T, where x is any amino acid except proline. Additionally, IgG contains one N-linked glycosylation site within the Fc. As such, we applied a liquid-chromatography-mass spectrometry (LC-MS) based approach to determine the site-specific glycosylation of ePGDM1400v9 IgA and IgG delivered as mRNA and produced by the macaque.

Analysis of mRNA delivered IgG demonstrated glycan processing that is typical for IgG. The three most abundant glycan compositions detected are within G0F (30%), G1F (32%) and G2F (11%) (Figure 3A and Table S3). The overall fucosylation was high, but not complete, with 81% of detected glycans containing fucose. In contrast to human-derived IgG, no N-acetylneuraminic acid was observed and instead 4% of glycans were modified with N-glycolylneuraminic acid (NeuGc) (Figure 3B). This monosaccharide is not found in humans as they lack the CMAH gene (which encodes cytidine monophosphate-N-acetylneuraminic acid hydroxylase). This enzyme is responsible for converting Neu5Ac (N-acetylneuraminic acid) to NeuGc. This observation highlights how glycosylation can diverge depending on the producer cell. A low level of under processed oligomannose-type glycans were also detected, which is also typical for IgG glycosylation.

**Fig. 3.**
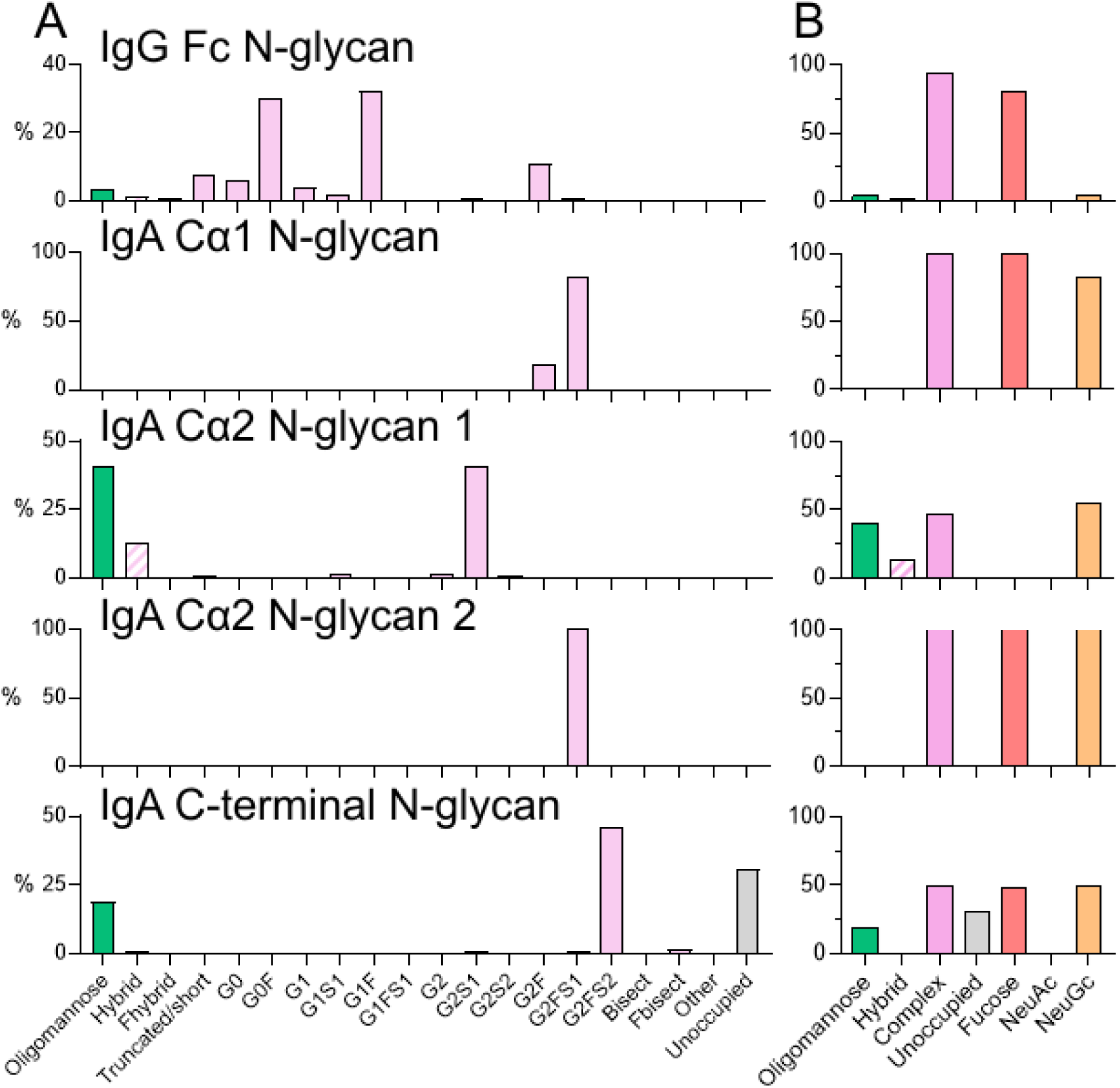
Site-specific glycan analysis of mRNA-derived ePGDM1400v9 IgG and IgA from transfused macaques. (A) Summary of the site-specific glycosylation of mRNA-derived ePGDM1400v9 IgA2. The relative proportion of different glycan types is summarized. The total proportion of glycans with compositions corresponding to oligomannose-type glycans is shown in green, hybrid-type glycans in hashed pink, complex-type glycans in pink. The proportion of potential N-linked glycan sites lacking glycan attachment (unoccupied) is shown in grey. As macaques contain NeuGc and no NeuAc was detected, categories denoted by “S” describe glycans containing NeuGc. (B) Summarized glycan categories for oligomannose-hybrid- and complex-type glycans. Additionally, the proportion of glycans containing one or more fucose monosaccharides is colored red, NeuAc (sialic acid) in purple and NeuGc (N-glycolylneuraminic acid) in orange. Glycan underoccupancy is also shown in grey.

The detected N-linked glycan compositions of IgA2 varied at the site-specific level, displaying a range of processing states including oligomannose-, hybrid- and complex-type glycans. Complex-type glycans were observed across all sites, ranging from 100% abundance on the Cα1 N-glycan site and Cα2 N-glycan site 2 to ∼50% on the Cα2 N-glycan site 1 (Figure 3 and Table S3). The majority of complex-type N-linked glycans were fucosylated and modified by NeuGc. Minimal levels of NeuAc were observed across all sites. N-linked glycans consisting of HexNAc(4)x were the predominant complex-type glycans observed across all sites, except for Cα2 N-glycan site 1 (Supplemental File - Glycopeptide analysis). Whilst linkage analysis was not possible with the selected LC-MS approach, N-linked glycans consisting of 4 N-acetylhexosamines typically correspond to biantennary glycans which comprise the G0, G1 and G2 categories typically used in IgG glycan nomenclature. Under-processed oligomannose-type glycans were detected at two sites, Ca2 glycan site 1 and the N-linked glycan site at the C-terminus. The presence of these glycans has been shown to enhance clearance (17). Previous analysis of IgA2 delivered mRNA to mice demonstrates a conservation of oligomannose-type glycans across species, and for both samples the overall abundance of oligomannose-type glycans is much lower from mRNA-derived IgA2 compared to recombinant protein, as observed previously (4). Finally, potential N-linked glycosylation sites lacking N-linked glycan attachment were observed at the C-terminal N-linked glycosylation site. As glycosylation occurs co-translationally, with the N-glycosylation attachment enzymes co-localizing with the ER-translocon, once translation is complete, N-linked glycosylation sites close to the C-terminus are often skipped as the polypeptide detaches from the ribosome complex (18).

## Discussion

Here, we demonstrate the expression of a mAb in NHPs via mRNA as both an IgG1 and an IgA2. Regardless of isotype, the mAb was expressed to detectable levels in the serum of all animals, and the mAb retained its ability to neutralize sensitive HIV pseudoviruses *in vitro*. Additionally, glycosylation patterns of mAbs produced by mRNA (both IgG1 and IgGA2) were more similar to native glycosylation patterns based on the observation of monosaccharide additions that are not usually seen in CHO/HEK-derived material (Figure 3). This study presents foundational data of proof-of-concept IgA expression in NHPs that could be further improved with subsequent designs and approaches.

All four NHPs that received mRNA-LNPs coding for ePGDM1400v9 IgA2 expressed detectable levels of the mAb, with peak serum levels of 1.4 – 5.4 µg/mL at 1 d post-transfusion, which was much lower than those observed for the IgG1 construct (Figure 1A). There are multiple potential explanations for this low expression level, which are not mutually exclusive. First, the IgA2 construct is transfected as three separate mRNA chains, each coding for a different polypeptide chain: the ePGDM1400v9 heavy chain, the ePGDM1400v9 light chain, and a J-chain to facilitate end-to-end dimerization of IgA2 monomers. Compared to the IgG1 construct, the extra polypeptide in this mix (J-chain) inherently reduces the molar quantity of heavy and light chain delivered per mg of mRNA-LNP. In conjunction with the heavy glycosylation typically present on IgA (19), this significantly increases the molecular complexity of the final product, which could be expected to reduce yields. The mRNA was formulated to provide a molar ratio of 1:2:1 and a mass ratio of 4:4:1 heavy:light:J chains, but this may not be the optimal ratio for producing IgA in NHPs. Additionally, ePGDM1400v9 may be a poor candidate for expression as an IgA: we have noticed as a recombinant protein produced by transient transfection of Expi293F cells, ePGDM1400v9 is not particularly well expressed or stable as an IgA2. Yields of the IgA2 construct in Expi293F cells were substantially lower than the IgG1 construct, and in PBS the IgA2 construct could not be concentrated above 0.22 mg/mL without precipitating whereas the IgG1 construct can be concentrated well over 50 mg/mL. Thus, attempting the experiment with antibodies more amenable to expression as an IgA and optimizing the molar ratio of IgA mRNA chains will likely improve *in situ* IgA expression levels in NHPs.

Two of the most attractive expected advantages of an IgA mAb over traditional IgG mAbs is the enhanced mucosal delivery, and the engagement of FcRαI (20). Unfortunately, the low expression levels of ePGDM1400v9 IgA2 observed preclude assessment of either of these features. Neutralization IC_50_ estimates suggest that, as expected, the mRNA-derived IgA2 in the serum and plasma was primarily monomeric (Figure 2E), which is not anticipated to traffic to mucosal sites. At 4 d post-transfusion, we were only able to detect ePGDM1400v9 IgA2 in the spleen and liver, where the mRNA-LNPs are expected to drive expression. However, our PK measurements suggest that by 4 d post-transfusion, serum ePGDM1400v9 IgA2 levels were probably only about 10% of peak levels, which were reached at 1 d post-transfusion (Figure 1A). Given that peak levels of mRNA-derived IgA2 were so low, and that a minority of total IgA2 produced may have been dimerized, the expected amount of IgA2 dimer in mucosal tissues at 4 d post-transfusion would likely be below our detection threshold even if it was successfully produced. Therefore, we are currently unable to distinguish between a failure to produce and traffic mRNA-derived IgA2 dimer to the mucosa and a failure to measure the low levels of dimer that were trafficked. Similarly, low expression levels mean that all available mRNA-derived IgA2 mAb was consumed for neutralization and glycosylation analysis, meaning that IgA-mediated effector functions will need to be assessed in future studies. Nonetheless, the data presented here show that it is possible to express a functional mAb as an IgA in NHPs, and this will inform future studies, which will need to achieve higher IgA expression levels and sample tissues at earlier timepoints post-transfusion to address these questions.

In contrast to the IgA2 data, mRNA-derived ePGDM1400v9 IgG1 expressed quite well, with PK that are comparable to those observed for recombinant ePGDM1400v9 in NHPs (IAVI, unpublished data) and the parent mAb PGDM1400 in humans (21). Comparison of the serum concentration and neutralization titers observed here to those reported previously with the recombinant PGDM1400 parent antibody in IgG1 format in murine challenge and human clinical studies suggests that the mRNA-derived mAb reached expression levels that should be protective (21, 22). In one animal, we even observed detectable serum levels of the mAb 70 days post-transfusion (Figure 1A). We also observed detectable ePGDM1400v9 IgG1 in all tissues (Figure 1D).

Altogether, these results demonstrate that mRNA-LNPs can generate functional ePGDM1400v9 IgG1 *in situ* at levels expected to be protective. This could have important implications for the prophylactic and therapeutic use of HIV bnAbs. HIV bnAbs are actively being explored as prophylactics against HIV infection, particularly in the context of mother-to-child transmission at birth. Though recent clinical studies suggest that an antibody cocktail with sufficient viral coverage could prevent HIV infection (23, 24), clinical implementation of such a therapeutic cocktail is currently cost-prohibitive. Since mRNA-LNPs bypass the expensive production of recombinant protein, this work provides an important proof-of-concept for mRNA delivery of HIV bnAbs in an NHP model. Future work can explore the delivery antibody cocktails by mRNA LNPs in NHPs and the protective efficacy of those cocktails against HIV/SIV infection in adult and infant macaques.

## Methods

### Recombinant antibody proteins

Recombinant ePGDM1400v9 IgA2 was purchased from GeneArt (ThermoFisher) whereby IgA2 protein was purified from transient transfection of a 4:4:1 ratio of HC to LC to JC by weight in Expi293 with CaptureSelect IgA Affinity matrix.

Recombinant ePGDM1400v9 IgG1 was produced via transient transfection of Expi293 cells using standard methods. To summarize, plasmids encoding for codon-optimized constructs of ePGDM1400v9 heavy and light chains were transfected at a 1:1 ratio by weight. Five days post-transfection, the antibody was purified by protein A/G affinity resin (Cytivia), and monodispersity was confirmed via size exclusion chromatography (SEC) using a Superdex S200 column (Cytivia). The ePGDM1400v9 Fab (with c-terminal His and FLAG tags) was performed using the same procedure, except that purification was performed using a Ni affinity column.

The anti-idiotype antibody aPGDM01 was raised against ePGDM1400v9 Fab using a commercial service. This antibody was validated by SPR, ELISA, and ECL (Figure S1A-S1E). It was produced as either a Fab or a Murine IgG2a via transient transfection into Expi293 cells as described above for ePGDM1400v9 IgG1.

### mRNA transfection

EXPI293 cells were transiently transfected with the corresponding mRNA encoding for ePGDM1400v9 IgG1 or IgA2 using *trans* IT-mRNA Transfection Kit (Mirus Bio LLC) per the manufacturer’s recommendations. For IgG, this required co-transfection of HC and LC at a 2:1 ratio by weight, whereas for IgA, the transfection consisted of HC, LC, and JC transfections at a 4:4:1 ratio by weight. EXPI293 cells were diluted to 1×10^6^ cells/mL with EXPI293 Expression Medium (ThermoFisher). Using a ratio of 1 μg mRNA per 1 mL of culture, mRNA was first added to Opti-MEM I SFM (Gibco), followed by addition of TransIT-mRNA transfection reagent (Mirus Bio) at a 1:2 ratio of mRNA to TransIT. The complex was allowed to form for 3 min before addition to the suspension culture. The quantity of antibody in the resultant supernatants was measured by ELISA.

### Generation of modified mRNA and LNPs

Sequence-optimized mRNA encoding functional IgA monoclonal antibodies were synthesized in vitro using an optimized T7 RNA polymerase-mediated transcription reaction with complete replacement of uridine by N1-methyl-pseudouridine (25). The reactions included a DNA template containing an open reading frame flanked by 5′ untranslated region (UTR) and 3′ UTR sequences and was terminated by an encoded polyA tail. Ligation reactions were performed using T4 RNA Ligase I (New England Biolabs) according to the manufacturers recommended conditions. mRNA was purified by dT affinity chromatography, buffer exchanged by ultrafiltration into 2 mM sodium citrate, sterile filtered, assayed for purity, and stored frozen at –20 °C for further use.

Lipid nanoparticle-formulated mRNA was produced through a modified ethanol-drop nanoprecipitation process as described previously (26). Briefly, ionizable, structural, helper, and polyethylene glycol lipids were mixed with mRNA (2:1 HC to LC by weight for IgG; 4:4:1 HC to LC to JC by weight for IgA) in acetate buffer, pH 5.0, at a ratio of 3:1 (lipids:mRNA). The mixture was neutralized with Tris-Cl, pH 7.5, sucrose was added as a cryoprotectant, and the final solution was sterile filtered. Vials were filled with formulated LNP and stored frozen at –70 °C until further use. The drug product underwent analytical characterization, which included the determination of particle size and polydispersity, encapsulation, mRNA purity, double stranded RNA content, osmolality, pH, endotoxin and bioburden, and the material was deemed acceptable for in vivo study.

### Research Animals and Study Design

Eight (6 male and 2 female) adult Indian-origin rhesus macaques were infused with liposomes containing mRNA expressing ePGDM1400v9-LS. Four macaques were infused with the mRNA encoding the antibody in an IgG1 format and four macaques were infused with the mRNA encoding the antibody in an IgA format. Blood and mucosal secretions were collected at multiple time points to collect pharmacokinetic data for each antibody. Rhesus macaques were housed at the Wisconsin National Primate Research Center (WNPRC). All procedures were approved by the University of Wisconsin-Madison College of Letters and Sciences and the Office of the Vice Chancellor for Research Institutional Animal Care and Use Committee (IACUC protocol number G006597). The animal facilities of the WNPRC are licensed by the US Department of Agriculture and accredited by AAALAC. Animals were monitored twice daily by veterinarians for any signs of disease, injury, pain, distress, or psychological abnormalities and treated as recommended by the veterinarian. Animals were euthanized at either 5- or 70-days post mRNA infusion. Animals were euthanized as recommended by the Panel on Euthanasia of the American Veterinary Medical Association by first administering general anesthesia (ketamine ≥15 mg/kg intramuscularly (IM)), then administering an intravenous (IV) overdose of sodium pentobarbital (≥ 50 mg/kg). After euthanasia tissues were taken at necropsy and either flash frozen, frozen into OCT blocks or embedded into FFPE blocks.

### LNP-mRNA infusion

LNP-mRNA infusion was performed under anesthesia (ketamine plus dexmedetomidine). Animals were prophylactically dosed intramuscularly with 1 mg/kg diphenhydramine 30 minutes prior to the infusion to prevent anaphylaxis. The animals were fitted with a catheter and the infusion of the mRNA was performed intravenously at a dose of 1 mg/kg over approximately a 1 h duration. Animals were closely monitored for reactions following the infusion and during anesthesia recovery. Blood was drawn longitudinally following infusion to monitor antibody levels.

### Sample collection, processing, and cryopreservation

Blood was collected from animals prior to the mRNA infusion, before and immediately after infusion, and then every 2 days through day 13 and then weekly for 5 weeks. Blood was collected into EDTA tubes and SST tubes and centrifuged twice to remove all cells and collect plasma and serum, respectively. Plasma and serum were stored at −80 °C.

Rectal and nasal secretions were sampled onto pre-weighed Weck-Cel sponges (Fisher Scientific) while the animals were under anesthesia (ketamine up to 7 mg/kg IM and up to 0.03 mg/kg dexmedetomidine IM, reversed with 0.3 mg/kg atipamezole (IV or IM)). Sponges were pre-moistened with sterile PBS and placed atraumatically into the rectal or nasal cavity for five minutes. Sponges were removed and placed into a sterile microcentrifuge tube. The post-collection swab was weighed and then the stick cut off, so the sponge fit inside the microcentrifuge tubes. Sponges were stored at −80 °C until tested.

### Tissue processing

NHP tissues were collected at necropsy and shipped to Charles River Laboratories for processing. Tissues pieces were weighed and cut to be approximately between 5 and 20 mg. Samples were added to pre-chilled Matrix D tubes (MP Biomedicals #6913050) and 200 uL of Tris lysing buffer (MSD #R60RTX) was added prior to placing the tubes on the FastPrep-24 homogenizer (MP Biomedical #116004500). For all tissues, the FastPrep-24 homogenizer was set to a speed of 6 meters/second for a time of 30 seconds. Following the first run, tubes were placed on wet ice for at least 5 minutes after which the FastPrep-24 homogenization was repeated 2 more times, with 5-minute incubation on ice in between. All samples were visually inspected for complete homogenization and then spun in a centrifuge at 20000 x g at 5 °C for 20 minutes. Supernatants were transferred to new tubes on wet ice and the spin was repeated. Following the final spin, supernatants were transferred to new tubes and stored at –80 °C until use. Collected tissues used for BioD analysis included Lung, vaginal, Rectal, Ascending Colon, Duodedum, Jejunum, Inguinal and teal LN, Spleen, Liver

### Ig extraction from NHP nasal and rectal wecks

The neutralization assay was performed on antibody extracted from nasal and rectal wecks. Briefly, each weck tubes received 150 µL of weck buffer (1xPBS, 10% normal goat serum and, for rectal wecks only, protease inhibitor) in which they incubated for 2 hours at RT. Each weck is then transfered to a new tube using forceps, and centrifugated at max speed for 5 min before being squeezed with tweezers, to extract as much remaining buffer as possible.

The recovered volume will be combined with the first extraction volume.

### Serum and tissue Ig concentration measurement

Serum and tissue monoclonal IgG1 and IgA2 were measured using MSD ECL. The assay (Figures S1D and S1E) was standardized using ePGDM1499v9-free NHP serum spiked with recombinant ePGDM1400v9 (either human IgG1 or human IgA2). 96 well MSD QuickPlex Standard plates (MSD Cat# L55XA) were coated in PBS with anti-human IgG CH2 domain (BioRad MCA5748G) for IgG and goat anti human kappa (Southern Biotech 2042-01) for IgA2. All coated plates were stored at 2-8 °C until use. Prior to use, coated plates were removed from 2-8 °C and allowed to warm to room temperature. Assay plates were washed 3 times with 300 µL per well of ELISA wash buffer (0.05% Tween 20 in TBS-T). Plates were blocked with 150 µL per well of 5% MSD Blocker A (MSD R93AA-1), sealed and incubated on a plate shaker (BoekelScientific 130000-2 microplate shaker) at room temperature for 2 h. Fresh standards and quality controls were thawed on the bench, briefly spun down and vortexed. ePGDM1400 IgG1 and IgA2 recombinant proteins served as standards for the respective assay. Standards for all serum ECL assays were diluted in heat inactivated normal rhesus macaque serum and for tissue ECL assays, standards were diluted in 5% Tris lysis buffer. For serum IgG1, standards ranged from 500 ng/mL to 1.95 ng/mL and for serum and tissue IgA2, standards ranged from 500 ng/mL to 0.98 ng/mL. Tissue IgG1 standards started at 300ng/mL and were diluted down to 0.0457 ng/mL. All standards and samples were diluted 1:10 (for serum ECL) and 1:20 (for tissue ECL) in low cross buffer (Boca Scientific 100-500) for a final volume of 100uL. Following blocking incubation, plates were washed 3x with ELISA washer buffer and 30 µL of diluted standards and samples were added to the plate in duplicate. Plates were sealed and incubated at RT on a plate shaker for 1 h. After primary incubation, plates were washed 3 times with ELISA wash buffer and 30 µL per well of secondary reagent was added. For both IgG1 and IgA2, a sulfotagged aPGDM01 diluted in 1% MSD Blocker A buffer was used (1:4000 dilution IgG1, 1:500 dilution IgA2). Plates were incubated at room temperature on a plate shaker for 1 hour. Following incubation, plates were washed 3 times with ELISA wash buffer and 150 µL of MSD Gold Read Buffer A was added to each well (MSD R92TG). All plates were read within 10 minutes. Sample concentrations were extrapolated from the standard curve using Watson 7.6.1 analysis software and a 5PL regression curve.

### Nasal and rectal weck Ig concentration measurement

Nasal and rectal wecks were stored frozen at –80 °C until processed. All wecks were inspected prior to processing for presence of feces and/or blood and this data was recorded. For both rectal and nasal wecks, there were two wecks per monkey per time point. Thawed rectal wecks were soaked in 150uL of PBS + 10% normal goat serum (ThermoFisher) and 1 protease inhibitor (Thermo Scientific) per 50mL of buffer for 2 h at room temperature. Thawed nasal wecks were soaked in 150 uL of PBS + 10% normal goat serum for 2 h at room temperature. Following incubation, wecks were taken out of buffer and placed with weck facing up in a new 1.5 mL eppendorf tube and spun at 14,000 rpm for 5 minutes to capture to dry out the weck and collect the fluid at the bottom of the tube. After centrifuging, tweezers were used to squeeze any remaining liquid out of the wecks and dispose of the weck. Centrifuge tubes again to collect all fluid at the bottom of the tube. Weck fluid was combined for the two wecks for each monkey to assess antibody levels in ELISA.

To assess levels of antibody in processes wecks, half-area 96 well plates (Corning Costar) were coated with 25 µL per well of 4 µg/mL of the anti-idiotype mAb aPGMD01 (as a murine IgG2a) overnight at 4 °C. Plates were washed using an automated plate washer (Biotek) with 0.05% PBS-Tween (PBS-T) and were subsequently blocked for 2 h with 50 µL per well of Superblock PBST (ThermoFisher). Following blocking, plates were washed again using an automated plate washer and incubated with weck samples at 4 °C overnight. Weck samples were added straight onto the plate and were not diluted. Purified recombinant ePGDM1400v9 IgG1 or IgA2 was used as standards on all plates. The next day, plates were washed with an automated plate washer (Biotek) and incubated with 25 µL per well either goat anti human IgA (alpha chain, cross adsorbed) HRP (Life Technologies, 1:10,000) or goat anti human H+L chain NHP cross-adsorbed HRP (American Qualex, 1:10,000) for 1 h at room temperature. Plates were subsequently washed and incubated with 25 µL per well of SureBlue TMB 1-C substrate (Fisher Scientific) for 5 minutes. The reaction was stopped with 25 µL per well of TMB Stop solution (SeraCare) and read at an absorbance of 450 nM on a SpectraMax ABS Microplate reader. Absolute quantities of human antibody in wecks were extrapolated in GraphPad Prism 9 using a standard curve that was generated with the appropriate purified isotype version of ePGDM1400v9.

### Neutralization Ig assay

Rhesus macaque sera or extracted Ig and control Igs were serially diluted in 96-well plates (Corning) and DMEM (Gibco) completed with 10% FBS (Gibco), 0.1 mg/mL Penicillin-Streptomycin (Gibco) and 2 mM L-Glutamine 2 mM (Gibco). 25 µL of each serum or antibody dilution were incubated with 25 µL of pseudotype virus supernatant for 1 h at 37 °C, prior addition of 50 µL per well TZMbl cells at 0.2 million cells/mL in suspension in DMEM complete (assay plate: Corning). The serum or antibody, virus and cells mix were incubated for 72 h at 37 °C prior medium aspiration, cell lysis and luciferase substrate addition (Luciferase Assay kit, Promega). Light intensity – reporter for infection - is measured with a luminometer (Biotek Synergy). The percentages of neutralization were calculated with excel and ID80 (dilution required to reach 80% neutralization against a given virus) were defined with GraphPad Prism.

### Surface Plasmon Resonance (SPR)

SPR measurements were performed on a Biacore 8K instrument at 25 °C. All experiments used HBS-EP+ (0.1M HEPES pH 7.6, 150 mM NaCl, 3 mM EDTA, 0.0005% (v/v) Surfactant P20) as a mobile phase at a flow rate of 30 µL/min. Anti-mouse IgG (Fc-specific) antibody (Cytivia Cat. No. BR100838) was immobilized on a Series S CM-3 sensor chip (Cytivia) using standard NHS/Edc coupling in accordance with the manufacturer’s instructions. A reference surface was generated using the same methods.

To validate that aPGMD01 bound ePGDM1400v9 in Fab, IgG1, and IgA2 formats, aPGMD01 was produced as mouse IgG2a mAbs and captured at 10 nM onto the anti-mouse IgG coated chips described above. A recombinant mouse IgG2a mAb that did not have detectable binding to ePGDM1400v9 by ELISA was used as a negative control. A concentration series of analyte (either recombinant ePGDM1400v9 Fab, human IgG1, or human IgA2) was injected across the antibody and control surfaces for 2 min followed by a 10 min dissociation step using a single-cycle method. Regeneration of the surface between injections of analyte was accomplished using 4-minute injections of 10 mM Glycine (pH 1.7). Data analysis was performed using BIAEvaluation software (Cytivia) and Prism (Graphpad).

### Purification of mRNA-produced ePGDM1400v9 from NHP sera/plasma

To purify mRNA-derived ePGDM1400v9 antibodies from NHP sera/plasma for glycan analysis, recombinant aPGDM01 was produced as a mouse IgG2a and coupled to CNBr resin (Cytivia Cat. No. 17043001) using the manufacturer’s protocol, except that PBS (pH 7.4) was used as coupling buffer.

Sera and/or plasma that had not been used for other experiments was pooled prior to applying it to the aPGDM01-coupled column. For purifying ePGDM1400v9 IgG, all available serum samples up to 42 d post-transfusion were pooled. For purifying ePGDM1400v9 IgA, plasma and sera from all animals up to 14 d for RMA3, up to 11 d for RMA4, and up to 5 d for all four NHPs based on sample availability and measured IgA from the PK measurements.

For the purifications, 250 µL of aPGDM01-coupled resin was washed with 10 mL of 1X PBS pH 7.4. Then, the pooled serum/plasma fractions were applied to the column by gravity flow. Next, the column was washed with 5 mL of salt wash buffer (0.1 M Tris pH 8.0, 500 mM NaCl), followed by a 5 mL detergent wash (1X PBS pH 7.4, 0.1% n-dodecyl-ß-D-maltoside (DDM)). A final wash of 5 mL 1X PBS pH 7.4 was then applied before eluting with 650 µL 0.1 M sodium citrate buffer, pH 3.0. The pH was neutralized by the addition of 350 µL 1 M Tris pH 8.0. The purified antibody was stored at 4 °C overnight. In the morning, the quality and quantity of the purified antibodies were confirmed by a Coomassie gel and a PGDM1400v9-specific ELISA (using plates coated with aPGDM01). The remaining preparations were then aliquoted and frozen at −20 °C until use.

### Glycosylation analysis of mRNA-produced ePGDM1400v9 purified from NHP plasma/sera

To analyze the site-specific glycosylation of mRNA-derived Ig, the pooled remaining protein was incubated for 1 h in 50 mM Tris/HCl, pH 8.0, containing 6 M of urea and 5 mM dithiothreitol (DTT). The sample was then incubated in the dark for 1 h with 20 mM iodoacetamide (IAA) to alkylate the protein. To remove the residual IAA, 20 mM DTT was added, and the sample was incubated for an additional 1 h period. Following buffer-exchange into 50 mM Tris/HCl, pH 8.0 using Vivaspin columns (10 kDa molecular weight cutoff). The proteins were then digested separately overnight at 37 °C with Trypsin (Promega) in a 1:30 (w/w) ratio.

Following digestion, a heated vacuum centrifuge set at 30 °C was used to dry down the peptides, which were then resuspended in 250 µL 0.1% TFA. Desalting and peptide enrichment was performed using an Oasis HLB desalting 96-well µElution plate (Waters) attached to a vacuum manifold. One well per digest was first equilibrated with 200 µL acetonitrile (ACN), followed by 200 µL 0.1% TFA. The peptides were loaded onto the elution plate at 1 mL/min, and the wells were then washed, first with 800 µL 0.1% TFA, followed by 200 µL of LC-MS grade H_2_O. The peptides were eluted by the addition of 70% ACN in 0.1% TFA and dried down again before being re-suspended in 0.1% formic acid. The peptides were combined and analyzed by nanoLC-ESI MS with a Vanquish Neo HPLC (Thermo Fisher Scientific) system coupled to an Orbitrap Eclipse mass spectrometer (Thermo Fisher Scientific). A μPAC™ Neo HPLC ColumnC18 column (75 µm x 110 cm) was used to separate the peptides. A trapping column (PepMap 100 C18 3 µM 75 µM x 2 cm) was used in line with the LC prior to separation with the analytical column. For LC separation, buffer A consisted of 0.1% formic acid and 80% acetonitrile in 0.1% formic acid. The LC conditions were as follows: 280-minute linear gradient consisting of 5-40% B in 0.1% formic acid over 240 minutes. The %B was then increased to 95% over 15 minutes and held for another 15 minutes before reducing the %B to 5%. The flow rate was set to 300 nL/min. The spray voltage was set to 2.5 kV. The ion transfer tube temperature was set to 275 °C. The scan range was 300−2000 m/z. Stepped HCD collision energy was set to 15, 25 and 45% and the MS2 for each energy was combined. Precursor and fragment detection were performed using an Orbitrap at a resolution MS1= 120,000. MS2= 30,000. The AGC target for MS1 was set to standard and injection time set to auto which involves the system setting the two parameters to maximize sensitivity while maintaining cycle time. Glycopeptide fragmentation data were extracted from the raw MS files using Byos (Version 4; Protein Metrics Inc). The glycopeptide fragmentation data were evaluated manually for each glycopeptide. The peptide was scored as true-positive when both the oxonium ions corresponding to the identified glycan and the correct b and γ fragment ions were observed.

The Protein Metrics 305 N-glycan library with sulphated glycans added manually, was used to search the MS data. The relative amounts of each glycan at each site in addition to the unoccupied proportion was determined by comparing the extracted chromatographic areas for different glycoforms with an identical peptide sequence. A 1% False discovery rate (FDR) was applied, and the precursor mass tolerance was set at 4 ppm, and 10 ppm for fragments. All charge sates for a single glycopeptide were summed. Glycans were categorized according to the composition detected. The assigned compositions can be found in Supplemental File - Glycopeptide analysis.

## Acknowledgments

We would like to acknowledge Finora Franck and Elizabeth Lampley for superb project management support. We would also like to thank Dr. Jordan Woehl for instruction and support in SPR. This study was made possible by the support of the American People through the United States Agency for International Development (USAID) and the US President’s Emergency Plan for AIDS Relief (PEPFAR) (grant No. AID-OAA-A-16-00032). The contents of this paper are the sole responsibility of the authors and do not necessarily reflect the views of USAID, PEPFAR, or the United States Government. The WNPRC is supported by NIH Grant P51OD011106. We thank the Scientific Protocol Implementation (SPI) team and Immunology Services (IS) team at the WNPRC for supporting the NHP work.

---

**Fig S1:**
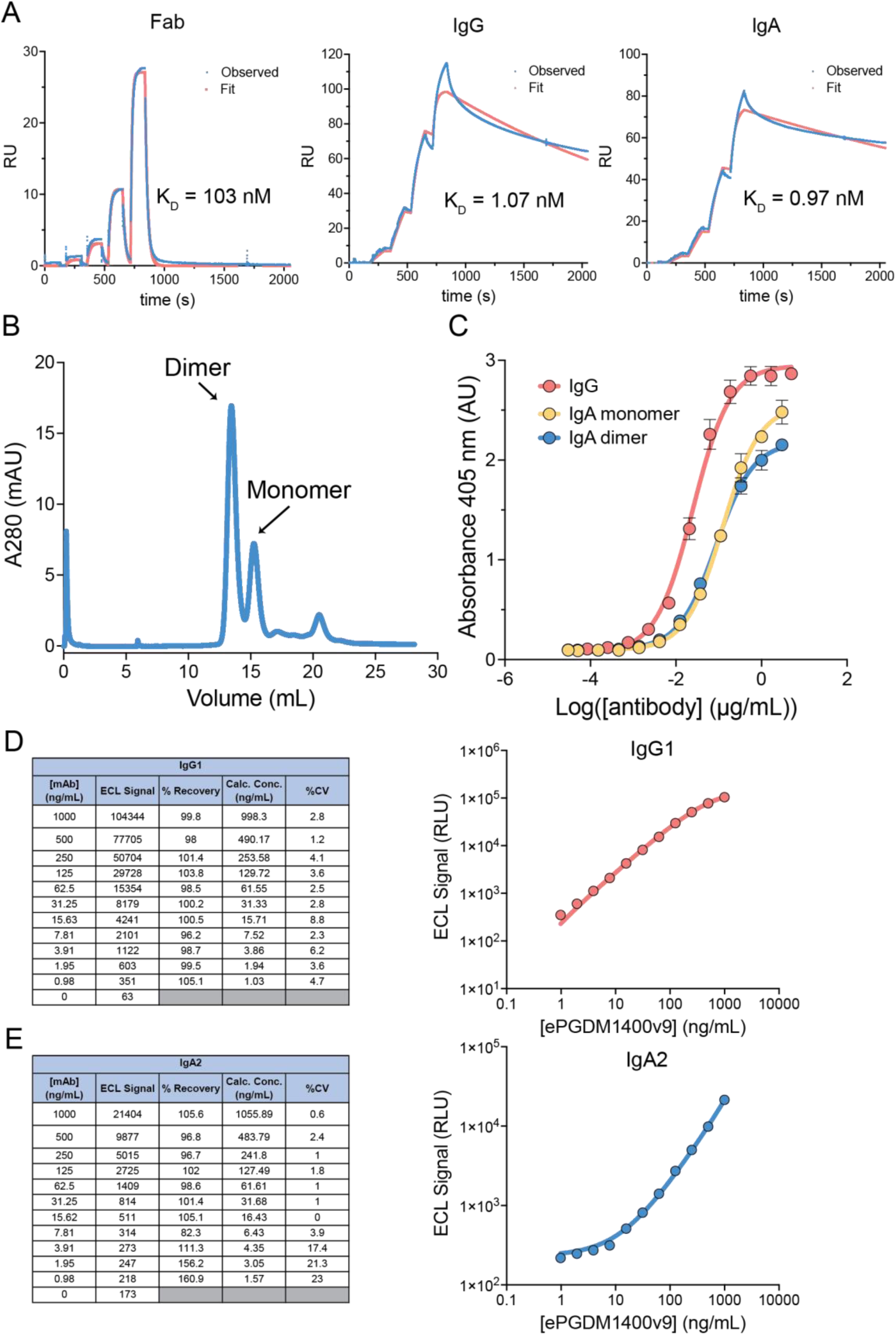
Anti-idiotype anti-ePGDM1400v9 mAb aPGDM01 enables ePGDM1400v9 IgG1 and IgA2 quantitation extrapolation via ECL. (A) Single-cycle SPR data using chips coated with aPGMD01 showing that the anti-idiotype mAb aPGMD01 binds ePGDM1400v9 as a Fab, IgG1, and IgA2. A negative control protein was run for each measurement and did not exhibit binding over background. (B) SEC trace of recombinant ePGDM1400v9 IgA2, showing distinct monomer and dimer peaks. Data acquired on a Superose 6 column in PBS (pH 7.4). (C) ELISA data showing binding of ePGDM1400v9 to plates coated with aPGDM01 mAb. Recombinant ePGDM1400v9 was diluted in PBS. Note that signal for IgG1 and IgA2 are not directly comparable, as different secondary antibodies were used. (D) Quantification of recombinant eGPDM1400v9 IgG1 diluted in non-mRNA infused NHP serum using the final aPGMD01-based ECL assay used to measure mRNA-derived ePGDM1400v9 IgG1 from NHPs infused with LNPs. (E) Quantification of recombinant eGPDM1400v9 IgA2 diluted in non-mRNA infused NHP serum using the final aPGMD01-based ECL assay used to measure mRNA-derived ePGDM1400v9 IgA2 from NHPs infused with LNPs.

**Fig. S2:**
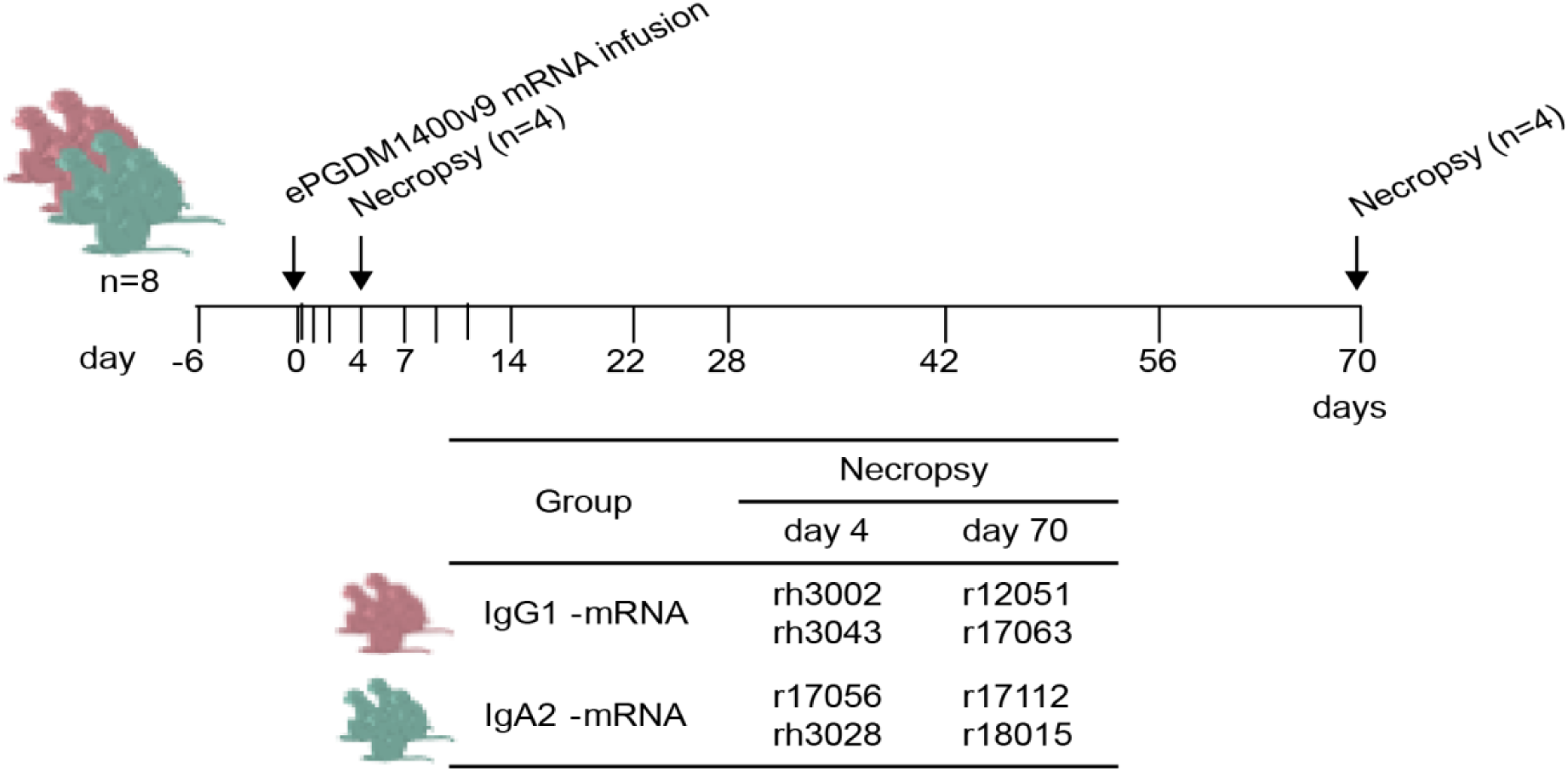
Experimental design for assessment of mRNA-delivered ePGDM1400v9 in IgG1 or IgA2 isotype biodistribution *in vivo*. A total of 8 rhesus macaques received ePGDM1400v9-IgG1 or ePGDM1400v9-IgA2 encoding mRNA at 1 mg/kg i.v. at day 0. Half (n=2) animals per groups were necropsied at day 4 and the second half at day 70. Plasma, Serum, nasal and rectal wecks were collected at −6 h, 0 h, and +6 h, and 1 d, 2 d, 4 d, and for the 4 remaining animals, also at 7 d, 9 d, 11 d, 14 d, 22 d, 28 d, 42 d, 56 d and 70 d.

**Table S1:**
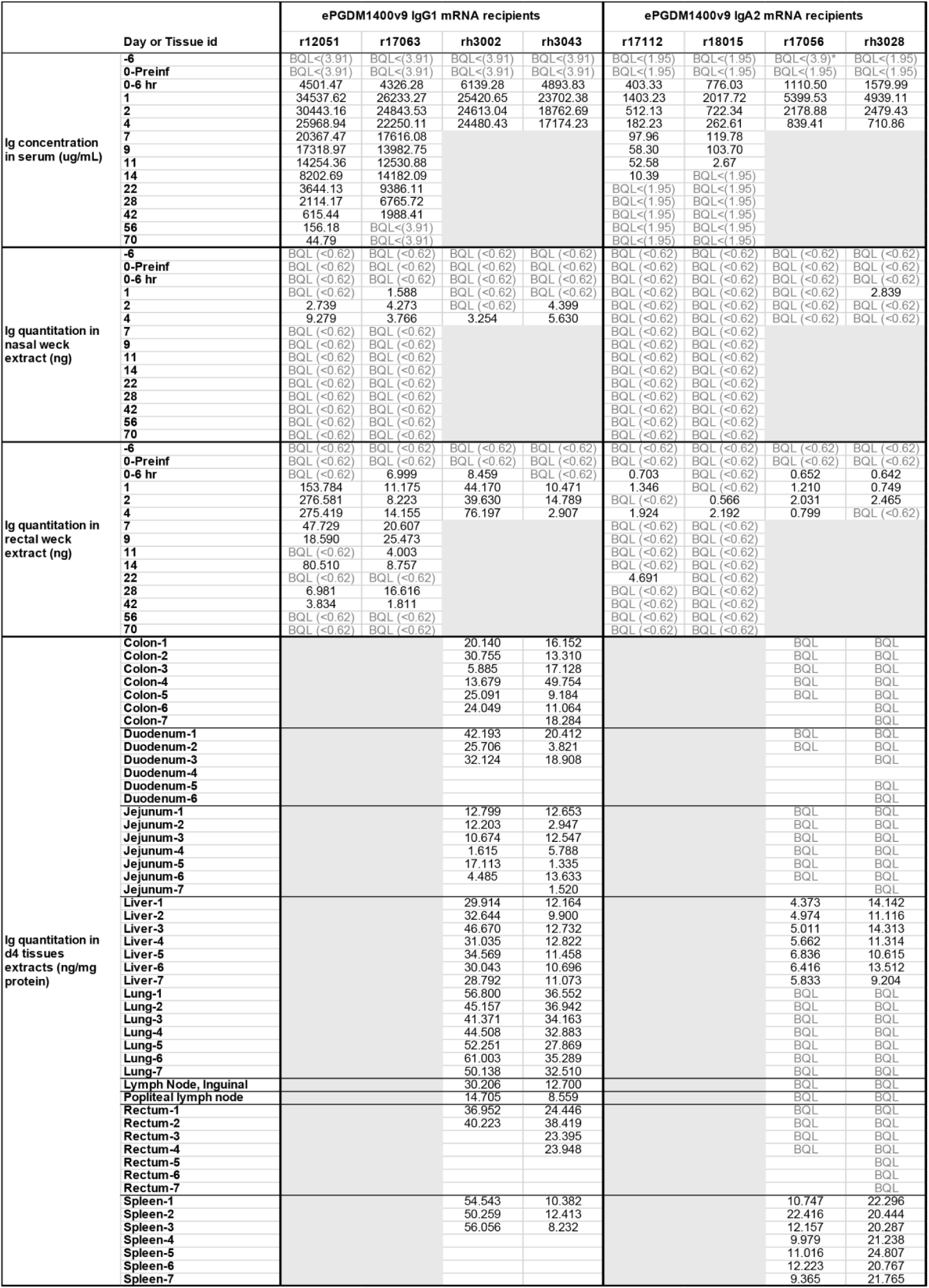
mRNA-delivered ePGDM1400v9 in IgG1 or IgA2 isotype quantitation in serum, nasal and rectal wecks and different tissues.

**Table S2:** Longitudinal neutralization ID_50_ of the NHP sera samples.

| Day or Tissue id |  | ePGDM1400v9 IgG1 mRNA recipients |  |  |  | ePGDM1400v9 IgA2 mRNA recipients |  |  |  |
| --- | --- | --- | --- | --- | --- | --- | --- | --- | --- |
|  |  | RM5 | RM7 | RM1 | RM3 | RM6 | RM8 | RM4 | RM2 |
| Serum neutralization ID50 against T250-4 (AG) | -6 | <20 | <20 | <20 | <20 | <20 | <20 | <20 | <20 |
|  | 0-Preinf | <20 | <20 | <20 | <20 | <20 | <20 | <20 | <20 |
|  | 0-6 hr | 5124 | 1045 | 706 | 4276 | 27 | 31 | 33 | 48 |
|  | 1 | 26318 | 6838 | 4449 | 17655 | 142 | 192 | 359 | 355 |
|  | 2 | 22611 | 9731 | 4458 | 15173 | 65 | 73 | 222 | 211 |
|  | 4 | 23021 | 9932 | 4900 | 14132 | 33 | 34 | 135 | 109 |
|  | 7 | 19746 | 5848 |  |  | <20 | <20 |  |  |
|  | 9 | 3522 | 3395 |  |  | <20 | <20 |  |  |
|  | 11 | 2960 | 2527 |  |  | <20 | <20 |  |  |
|  | 14 | 1686 | 3014 |  |  | <20 | <20 |  |  |
|  | 22 | 900 | 1744 |  |  | <20 | <20 |  |  |
|  | 28 | 528 | 1577 |  |  | <20 | <20 |  |  |
|  | 42 | 111 | 644 |  |  | <20 | 33 |  |  |
|  | 56 | 22 | <20 |  |  | <20 | <20 |  |  |
|  | 70 | 20 | <20 |  |  | <20 | <20 |  |  |
| Serum neutralization ID50 against CE1176 (C) | -6 | <20 | <20 | <20 | <20 | 144 | <20 | <20 | <20 |
|  | 0-Preinf | <20 | <20 | <20 | <20 | 96 | <20 | <20 | <20 |
|  | 0-6 hr | <20 | 54 | 30 | 55 | 115 | 23 | 21 | 34 |
|  | 1 | 527 | 520 | 385 | 279 | 233 | 123 | 286 | 259 |
|  | 2 | 449 | 597 | 381 | 328 | 174 | 48 | 129 | 119 |
|  | 4 | 62 | 465 | 218 | 321 | 143 | <20 | 43 | 42 |
|  | 7 | 84 | 259 |  |  | 131 | <20 |  |  |
|  | 9 | 131 | 155 |  |  | 157 | <20 |  |  |
|  | 11 | 100 | 260 |  |  | 168 | <20 |  |  |
|  | 14 | 122 | 167 |  |  | 155 | <20 |  |  |
|  | 22 | <20 | 89 |  |  | 156 | <20 |  |  |
|  | 28 | <20 | 48 |  |  | 156 | <20 |  |  |
|  | 42 | <20 | <20 |  |  | 180 | <20 |  |  |
|  | 56 | <20 | <20 |  |  | 158 | <20 |  |  |
|  | 70 | <20 | <20 |  |  | <20 | <20 |  |  |
| Serum neutralization ID50 against 6535 (B) | -6 | <20 | <20 | <20 | <20 | 32 | <20 | <20 | <20 |
|  | 0-Preinf | <20 | <20 | <20 | <20 | 21 | <20 | <20 | <20 |
|  | 0-6 hr | <20 | <20 | <20 | <20 | 28 | <20 | <20 | <20 |
|  | 1 | <20 | <20 | <20 | <20 | 62 | <20 | <20 | <20 |
|  | 2 | <20 | <20 | <20 | <20 | 31 | <20 | <20 | <20 |
|  | 4 | <20 | <20 |  |  | 28 | <20 | <20 | <20 |
|  | 7 | <20 | <20 |  |  | <20 | <20 |  |  |
|  | 9 | <20 | <20 |  |  | 22 | <20 |  |  |
|  | 11 | <20 | <20 |  |  | 25 | <20 |  |  |
|  | 14 | <20 | <20 |  |  | <20 | <20 |  |  |
|  | 22 | <20 | <20 |  |  | <20 | <20 |  |  |
|  | 28 | <20 | <20 |  |  | <20 | <20 |  |  |
|  | 42 | <20 | <20 |  |  | 23 | <20 |  |  |
|  | 56 | <20 | <20 |  |  | <20 | <20 |  |  |
|  | 70 | <20 | <20 |  |  | <20 | <20 |  |  |
| Serum neutralization ID50 against ML | -6 | <20 | <20 | <20 | <20 | <20 | <20 | <20 | <20 |
|  | 0-Preinf | <20 | <20 | <20 | <20 | <20 | <20 | <20 | <20 |
|  | 0-6 hr | <20 | <20 | <20 | <20 | <20 | <20 | <20 | <20 |
|  | 1 | <20 | <20 | <20 | <20 | <20 | <20 | <20 | <20 |
|  | 2 | <20 | <20 | <20 | <20 | <20 | <20 | <20 | <20 |
|  | 4 | <20 | <20 | <20 | <20 | <20 | <20 | <20 | <20 |
|  | 7 | <20 | <20 |  |  | <20 | <20 |  |  |
|  | 9 | <20 | <20 |  |  | <20 | <20 |  |  |
|  | 11 | <20 | <20 |  |  | <20 | <20 |  |  |
|  | 14 | <20 | <20 |  |  | <20 | <20 |  |  |
|  | 22 | <20 | <20 |  |  | <20 | <20 |  |  |
|  | 28 | <20 | <20 |  |  | <20 | <20 |  |  |
|  | 42 | <20 | <20 |  |  | <20 | <20 |  |  |
|  | 56 | <20 | <20 |  |  | <20 | <20 |  |  |
|  | 70 | <20 | <20 |  |  | <20 | <20 |  |  |

**Table S3:** Longitudinal neutralization ID_50_ of the NHP nasal weck samples.

| Day or Tissue id |  | ePGDM1400v9 IgG1 mRNA recipients |  |  |  | ePGDM1400v9 IgA2 mRNA recipients |  |  |  |
| --- | --- | --- | --- | --- | --- | --- | --- | --- | --- |
|  |  | RM5 | RM7 | RM1 | RM3 | RM6 | RM8 | RM4 | RM2 |
| Extracted Ig from nasal weck neutralization ID50 against T250-4 (AG) | -6 |  |  |  |  |  |  |  |  |
|  | 0-Preinf | <50 | <50 | <50 | <50 | <50 | <50 | <50 | <50 |
|  | 0-6 hr | <50 | <50 | <50 | <50 | <50 | <50 | <50 | <50 |
|  | 1 | nd | nd | <50 | nd | 814 | 777 | nd | <50 |
|  | 2 | <50 | <50 | <50 | <50 | <50 | <50 | <50 | <50* |
|  | 4 | <50 | <50 | <50 | <50 | <50 | <50 | nd | <50 |
|  | 7 | nd | <50 |  |  | <50 | <50 |  |  |
|  | 9 | <50 | nd |  |  | <50 | <50 |  |  |
|  | 11 | <50 | <50 |  |  | <50 | <50 |  |  |
|  | 14 | nd | nd |  |  | <50 | <50 |  |  |
|  | 22 | <50 | <50 |  |  | <50 | <50 |  |  |
|  | 28 | 56 | nd |  |  | 365 | nd |  |  |
|  | 42 | 326 | nd |  |  | nd | nd |  |  |
|  | 56 | <50 | <50 |  |  | <50 | <50 |  |  |
|  | 70 | 27 | <50 |  |  | <50 | nd |  |  |
| Extracted Ig from nasal weck neutralization ID50 against CE1176 (C) | -6 |  |  |  |  |  |  |  |  |
|  | 0-Preinf | <50 | <50 | <50 | <50 | <50 | <50 | <50 | <50 |
|  | 0-6 hr | <50 | <50 | <50 | <50 | <50 | <50 | <50 | <50 |
|  | 1 | nd | nd | <50 | nd | 2211 | 2667 | nd | <50 |
|  | 2 | <50 | <50 | <50 | <50 | <50 | <50 | <50 | <50* |
|  | 4 | <50 | <50 | <50 | <50 | <50 | <50 | nd | <50 |
|  | 7 | nd | <50 |  |  | <50 | <50 |  |  |
|  | 9 | <50 | nd |  |  | <50 | <50 |  |  |
|  | 11 | <50 | <50 |  |  | <50 | <50 |  |  |
|  | 14 | nd | nd |  |  | <50 | <50 |  |  |
|  | 22 | <50 | <50 |  |  | <50 | <50 |  |  |
|  | 28 | <50 | nd |  |  | 371 | nd |  |  |
|  | 42 | 73 | nd |  |  | nd | nd |  |  |
|  | 56 | <50 | <50 |  |  | <50 | <50 |  |  |
|  | 70 | <50 | <50 |  |  | <50 | nd |  |  |
| Extracted Ig from nasal weck neutralization ID50 against 6535 (B) | -6 |  |  |  |  |  |  |  |  |
|  | 0-Preinf | 666 | <50 | <50 | <50 | <50 | <50 | <50 | <50 |
|  | 0-6 hr | 1279 | <50 | <50 | <50 | <50 | <50 | <50 | <50 |
|  | 1 | nd | nd | <50 | nd | 1518 | 4087 | nd | <50 |
|  | 2 | 12114 | <50 | <50 | <50 | <50 | <50 | <50 | <50* |
|  | 4 | 860 | <50 | <50 | <50 | <50 | <50 | nd | <50 |
|  | 7 | nd | <50 |  |  | <50 | 63 |  |  |
|  | 9 | 1088 | nd |  |  | <50 | <50 |  |  |
|  | 11 | 2074 | <50 |  |  | <50 | <50 |  |  |
|  | 14 | nd | nd |  |  | <50 | <50 |  |  |
|  | 22 | 5640 | <50 |  |  | <50 | 64 |  |  |
|  | 28 | 633 | nd |  |  | 8197 | nd |  |  |
|  | 42 | 2960 | nd |  |  | nd | nd |  |  |
|  | 56 | 1313 | 67 |  |  | 2534 | <50 |  |  |
|  | 70 | 1183 | <50 |  |  | <50 | nd |  |  |
| Extracted Ig from nasal weck neutralization ID50 against ML | -6 |  |  |  |  |  |  |  |  |
|  | 0-Preinf | <50 | <50 | <50 | <50 | <50 | <50 | <50 | <50 |
|  | 0-6 hr | <50 | <50 | <50 | <50 | <50 | <50 | <50 | <50 |
|  | 1 | nd | nd | <50 | nd | <50 | 1467 | nd | <50 |
|  | 2 | <50 | <50 | <50 | <50 | <50 | <50 | <50 | <50* |
|  | 4 | <50 | <50 | <50 | <50 | <50 | <50 | nd | <50 |
|  | 7 | nd | <50 |  |  | <50 | <50 |  |  |
|  | 9 | <50 | nd |  |  | <50 | <50 |  |  |
|  | 11 | <50 | <50 |  |  | <50 | <50 |  |  |
|  | 14 | nd | nd |  |  | <50 | <50 |  |  |
|  | 22 | <50 | <50 |  |  | <50 | 33 |  |  |
|  | 28 | <50 | nd |  |  | 490 | nd |  |  |
|  | 42 | 73 | nd |  |  | nd | nd |  |  |
|  | 56 | <50 | <50 |  |  | 94 | <50 |  |  |
|  | 70 | <50 | <50 |  |  | <50 | nd |  |  |

\* presence of blood on sample nd = not determined (volume too low)

**Table S4:**
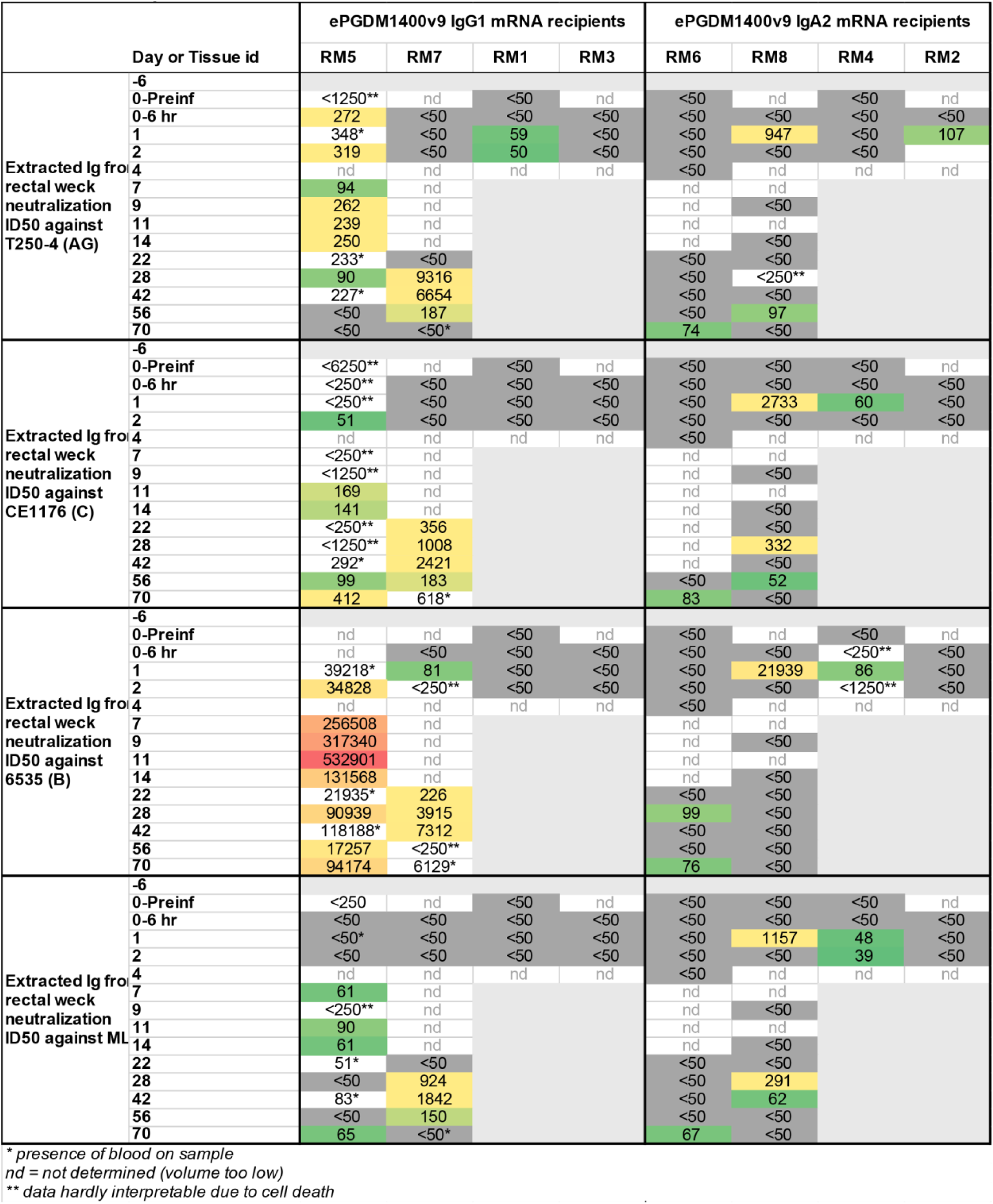
Longitudinal neutralization ID_50_ of the NHP rectal weck samples.

**Table S5:**
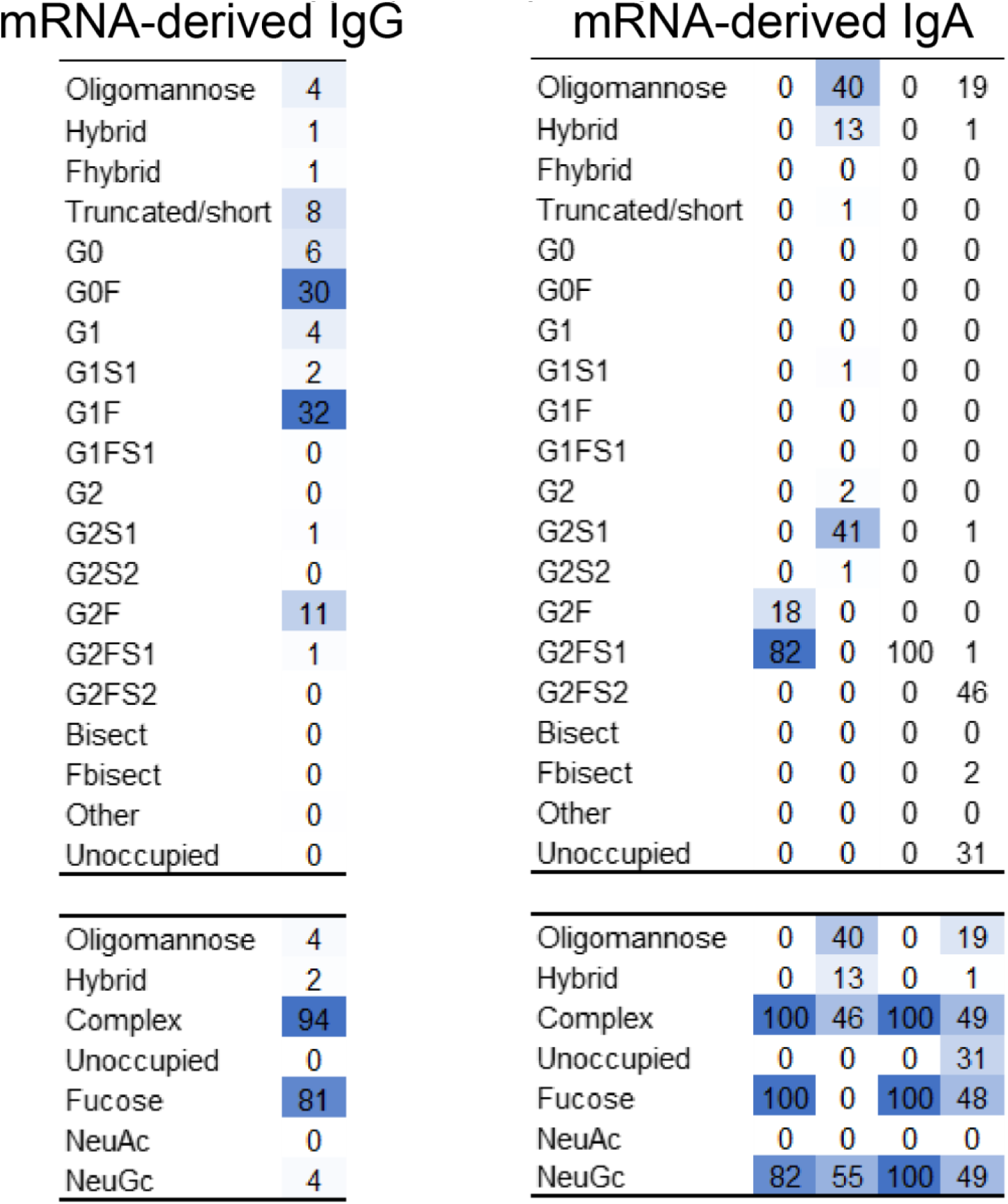
Site-specific glycosylation of IgG and IgA.

|  |  |
| --- | --- |
| Oligomannose | 4 |
| Hybrid | 1 |
| Fhybrid | 1 |
| Truncated/short | 8 |
| G0 | 6 |
| G0F | 30 |
| G1 | 4 |
| G1S1 | 2 |
| G1F | 32 |
| G1FS1 | 0 |
| G2 | 0 |
| G2S1 | 1 |
| G2S2 | 0 |
| G2F | 11 |
| G2FS1 | 1 |
| G2FS2 | 0 |
| Bisect | 0 |
| Fbisect | 0 |
| Other | 0 |
| Unoccupied | 0 |
| Oligomannose | 4 |
| Hybrid | 2 |
| Complex | 94 |
| Unoccupied | 0 |
| Fucose | 81 |
| NeuAc | 0 |
| NeuGc | 4 |

|  |  |  |  |  |
| --- | --- | --- | --- | --- |
| Oligomannose | 0 | 40 | 0 | 19 |
| Hybrid | 0 | 13 | 0 | 1 |
| Fhybrid | 0 | 0 | 0 | 0 |
| Truncated/short | 0 | 1 | 0 | 0 |
| G0 | 0 | 0 | 0 | 0 |
| G0F | 0 | 0 | 0 | 0 |
| G1 | 0 | 0 | 0 | 0 |
| G1S1 | 0 | 1 | 0 | 0 |
| G1F | 0 | 0 | 0 | 0 |
| G1FS1 | 0 | 0 | 0 | 0 |
| G2 | 0 | 2 | 0 | 0 |
| G2S1 | 0 | 41 | 0 | 1 |
| G2S2 | 0 | 1 | 0 | 0 |
| G2F | 18 | 0 | 0 | 0 |
| G2FS1 | 82 | 0 | 100 | 1 |
| G2FS2 | 0 | 0 | 0 | 46 |
| Bisect | 0 | 0 | 0 | 0 |
| Fbisect | 0 | 0 | 0 | 2 |
| Other | 0 | 0 | 0 | 0 |
| Unoccupied | 0 | 0 | 0 | 31 |
| Oligomannose | 0 | 40 | 0 | 19 |
| Hybrid | 0 | 13 | 0 | 1 |
| Complex | 100 | 46 | 100 | 49 |
| Unoccupied | 0 | 0 | 0 | 31 |
| Fucose | 100 | 0 | 100 | 48 |
| NeuAc | 0 | 0 | 0 | 0 |
| NeuGc | 82 | 55 | 100 | 49 |

## Notes

### Competing Interest Statement

Romy Rouzeau and Hayden Schmidt are employees of IAVI, which holds patents on ePGDM1400v9 and is advancing this antibody as a clinical candidate for HIV prophylaxis. Andrea Carfi is an employee of Moderna Tx, which owns the mRNA-LNP platform used here. Cailin Deal, Obadiah Plante, and Andrea Carfi have stock in Moderna Tx. Devin Sok is listed as an inventor on the patent for the parent antibody of ePGDM1400v9, PGDM1400 (US patent USRE50268E1).

